# Comparative transcriptomics of albino and warningly coloured caterpillars

**DOI:** 10.1101/336636

**Authors:** Juan A. Galarza

## Abstract

Colouration is perhaps one of the most prominent adaptations for survival and reproduction of most taxa. Colouration is of particular importance for aposematic species, which rely on their colouring and patterning to act as a warning signal against predators. Most research has focused on the evolution of warning colouration by natural selection. However, little information is available for colour mutants of aposematic species, particularly at the genomic level. Here I compare the transcriptomes of albino mutant caterpillars of the wood tiger moth (*Arctia plantaginis*) to those of their full-sibs having their distinctive orange-black warning colouration. The results showed >300 differentially expressed genes transcriptome-wide. Genes involved in the immune system, structural constituents of cuticle, and peptidase activity were mostly down-regulated in the albino larvae. Surprisingly, higher expression was observed in core melanin genes from albino larvae, suggesting that melanin synthesis may be disrupted in terminal ends of the pathway during its final conversion. I further identified 25 novel genes uniquely expressed in the albino larvae. Functional annotation showed that these genes are involved in nucleotide biosynthesis, copper ion transmembrane transport, and nucleic acid binding. Taken together, these results suggest that caterpillar albinism may not be due to a depletion of melanin precursor genes. In contrast, the albino condition may result from the combination of faulty melanin conversion late in its synthesis and structural deficiencies in the cuticle preventing its deposition.

## Introduction

Colouration is perhaps one of the most prominent adaptations for survival and reproduction of most taxa. It serves a variety of functions such as concealment for protection or ambuscading, intra- or inter-specific communication for sexual signalling or advertisement, and regulation of physiological processes such as body temperature (Cott 1940). Colouration can be produced by both, physical (i.e. structural) and chemical (i.e. pigments) means and it can change with ontogeny and/or seasonality. Its research goes beyond the evolutionary/ecological fields by influencing breakthroughs in the design of new materials and technologies (Caro, et al. 2017).

Colouration is of particular importance for aposematic species, which rely on their colouring and patterning to act as a warning signal advertising unpalatability to predators (Rowe and Guilford 2000). Such anti-predator adaptation is widespread across plants and animals (Komarek 1998; Mappes, et al. 2005). Most research has focused on the evolution of warning colouration by natural selection where its effectiveness is relative to its intra-inter-specific frequency, and in the context of the predator community (Mappes, et al. 1999; Merilaita and Kaitala 2002; Mochida 2011; Nokelainen, et al. 2014; Papaj and Newsom 2005). At the genomic level, aposematic colouration and patterning have been extensively study, particularly in Lepidoptera (Brower 2013; Dasmahapatra, et al. 2012; Davey, et al. 2016; Mallet 2010; Zhan, et al. 2014). However, almost no genomic information is available for colour mutants in wild populations. This is mainly due to the difficulty of obtaining samples given their very low expected frequencies.

Colour mutants provide a rare excellent opportunity to confirm suggested pigmentation mechanisms, and to gain insight on alternative or complementary mechanisms involved in warning colouration and patterning. Mutations that have a major effect, such as a complete absence of pigmentation (i.e. albinism) can provide valuable information. Albinism and leucism (i.e. partial absence of pigmentation) are best studied in humans and other mammals (Crawford, et al. 2017; de Vasconcelos, et al. 2017; Martinez-Garcia and Montoliu 2013; Rothammer, et al. 2017). Nonetheless, such conditions can be found across taxa including fish (Clark 2002; Nobile, et al. 2016), reptiles (Krecsák 2008; Mitchell and Church 2002) and plants (Gettys and Wofford 2007; Us-Camas, et al. 2017) However, little information is available for aposematic species.

Here I compare the transcriptomes of albino mutant caterpillars of the wood tiger moth (*Arctia plantaginis*) to those of their normal coloured siblings having their orange-black warning colouration. I evaluate transcriptome-wide gene expression differences, identify uniquely expressed genes in each condition, and analyse candidate genes from the melanin biosynthesis pathway. A comprehensive annotation of the transcriptomes of both conditions as well as of the differentially expressed and unique genes is provided. These results represent a thorough characterisation of genes and processes associated with the lack of pigmentation in Lepidoptera and are discussed in the context of aposematic strategies.

## Materials and Methods

### Study Species

The aposematic wood tiger moth is widely distributed throughout the Holarctic. It has one generation per year and adults are sexually dimorphic. Female hindwing colouration varies continuously from orange-red, whereas male hindwing colouration varies within populations from yellow-red in the Caucasus, yellow-white in Europe and Siberia, and black-white in North America and Northern Asia (Hegna, et al. 2015). Caterpillars however, are dichromatic displaying an orange patch against an otherwise dark body. The patch is variable in size and functions as a warning signal against avian predators (Lindstedt, et al. 2008).

Wood tiger moths have been reared at the University of Jyväskylä, Finland since 2009. The stock is dynamic (i.e. not permanent selection lines) being supplemented every year with wild-caught adults from which an average of 15000 larvae/year are produced. Albino larvae are rare, with only 8 cases documented showing completely white skin, hairs, and eyes (supplementary material S8). In human nomenclature, these would be referred to as ocolcutaneous (OCA) albinism (Oetting and King 1999). Albino larvae usually die within a few days or, in some cases, regain their warning colouration in the next moult, suggesting that this condition can be lethal or transient.

### Sample Collection and RNA-Sequencing

In this study, one albino larva was collected in summer 2011 and another in summer 2014. Both larvae were F1 from wild-caught parents. En each case, one of their normal coloured full-sibs was collected as well for comparative purposes. All larvae originated from Finnish populations and were in the fourth or fifth instar. They were fed with wild dandelion (*Taraxacum spp.*) and reared under natural light conditions with an average temperature of 25°C during the day and 15 - 20°C at night. For RNA extraction, larvae were submerged in RNAlater stabilizing solution (Qiagen, Valencia, US.A.) and kept at −20C° until extraction. Total RNA was extracted using RNeasy Mini Kit (Qiagen) according to manufacturer’s instructions with additional TriReagent (MRC, Inc.) and DNase (Qiagen, Valencia, US.A.) treatments. The quality and quantity of total RNA was inspected in a BioAnalyzer 2100 using RNA 6000 Nano Kit (Agilent). Subsequently, mRNA was isolated by means of two isolation cycles using Dynabeads mRNA purification kit (Ambion®) and quantified using RNA 6000 Pico Kit in a BioAnalyzer 2100 (Agilent). Pair-end (2 × 100pb) cDNA libraries were constructed for each larva according to Illumina’s TruSeq Stranded HT protocol. The libraries were individually indexed and sequenced in a HiScanSQ sequencer at the DNA sequencing and genomics laboratory of the Institute of Biotechnology at the University of Helsinki, Finland.

### Reads Processing and Mapping

The quality of the raw reads from the libraries was first inspected with FastQC (http://www.bioinformatics.babraham.ac.uk/pro-jects/fastqc/) and summarised using MultiQC v0.8 (Ewels, et al. 2016). Based on this initial quality check, the FASTX toolkit (http://hannonlab.cshl.edu/fastx_toolkit/) was used to remove low quality bases and sequencing artefacts. Bases with a Phred quality score of less than 25 were filter out, and reads shorter than 85 bases after trimming were removed. The remaining high-quality clean pair-end reads were then sorted and synchronized using custom scripts.

### Transcriptome Assembly

The high quality reads from the coloured larvae were pooled together and used to construct a *de novo* transcriptome assembly (Col_Trans hereafter). The same was done with the high quality reads from the albino larvae (Al_Trans hereafter). The transcriptomes were assembled according to Galarza, et al. (2017). Briefly, for each transcriptome, the software Trinity v.r2013-02-25 (Grabherr, et al. 2011) was first used to create an initial assembly with default K-mer =25 parameter. This recovers most of the full-length transcripts. Any unassembled reads were identified by mapping all reads back to the initial assembly using bowtie2 v. 2.2.5 (Langmead and Salzberg 2012). The unassembled reads were then used to construct two further assemblies with K-mer =21 and K-mer =29. Finally, a consensus transcriptome was constructed by combining the three assemblies through scaffolding using CAP3 (Huang and Madan 1999). Trinity was run using 16 CPUs with a memory cap of 200 GB and a minimum contig length of 500 bp. The CAP3 software was run with default parameters setting minimum overlap between two contigs of at least 100 bp with a 95% sequence similarity for obtaining of the final unigenes. Duplicate unigenes were removed using BBTools (https://jgi.doe.gov/data-and-tools/bbtools/bb-tools-user-guide/). All raw reads and both assemblies can be found at National Center for Biotechnology Information (NCBI) under project number PRJNA449279 with accession GGLW00000000.

### Transcriptome Validation and Annotation

The two transcriptomes were first aligned (BLASTn, e-value ≤ 10^−5^) against the curated ribosomal database SILVA (Quast, et al. 2013) (SILVA_132_SSU_Nr99_tax & SILVA_132_LSUR_Ref_tax Release 11-12-2017) to filter out possible contaminants (i.e. sequences from archaea, bacteria, and eukaryote domains). Transcriptome completeness was then evaluated with BUSCO v.3 (Simão, et al. 2015) using the insecta lineage. The transcriptomes were then aligned (BLASTx) (Altschul, et al. 1997) against a non-redundant protein databases (nr) (NCBI; last updated 15-06-2017) and the Swiss-Prot (last updated 25– 06-2017) to retrieve functional annotation. After blasting, all hits that showed < 70% amino acid identity, alignment length of < 100 bp amino acid length, and e-value ≤ 10-5 were removed. Gene ontology terms (GO) and information of protein family was obtained using Blast2Go v.4.0 (Conesa, et al. 2005).

### Differential Gene Expression

To investigate gene expression differences between the two larval conditions, the high quality reads from the four samples were first mapped to the wood tiger moth’s reference transcriptome (Galarza, et al. 2017) using bowtie2 v. 2.2.5 (Langmead and Salzberg 2012). The number of mapped reads for each sample was counted using SAMtools v.1.3.1 (Li 2009) and merged into a count matrix which was then normalised by transcript length and sequencing depth. Here, for each transcript, the number of mapped (Mr) reads was divided by the total number of reads (Tr), multiplied by transcript length (Tl) scaled by a factor of a million (i.e. (Mr/Tr)*^10^9^). This procedure returns normalized counts as transcripts per every million reads sequenced (TPM), in which the sum of all TPMs in each sample is the same, thus allowing a direct comparison of normalized expression values across samples and conditions. The R package edgeR (Robinson, et al. 2010) was used to test for differential gene expression setting a *P*-value cut-off threshold of 0.05, with a Benjamin and Hochberg correction (Benjamini and Hochberg 1995) for multiple testing. Subsequently, a functional annotation of the differentially expressed genes between conditions was obtained by blasting (BLASTx) the up- and down-regulated genes against non-redundant protein databases (nr) (NCBI; last updated 15-06-2017) the Swiss-Prot (last updated 25–06-2017). All hits that showed < 70% amino acid identity, alignment length of < 100 bp, and e-value ≤ 10-5 were excluded and the gene ontology terms (GO) and information of protein family was obtained using Blast2Go v.4.0 (Conesa, et al. 2005).

### Gene Expression Validation

To validate gene expression results from RNA-seq data, a subset of 6 differentially expressed gene transcripts was evaluated through quantitative PCR (qPCR). The expression of three gene transcripts involved in larval cuticular processes and three other non-annotated random gene transcripts, were selected for qPCR comparison to the RNA-seq data. Exon-intron boundaries were first identified by aligning these gene transcripts to genomic data previously obtained through 454 (Life Sciences) pyrosequencing using Mummer v.3.23 (Kurtz, et al. 2004). Bridging primers were then designed using Primer3 v. 4.0.0 (Untergasser, et al. 2012). As a normalization control (i.e. housekeeping gene), I selected one transcript from the RNA-seq data, which showed a uniform expression level within and between the two larval conditions. The software Normfinder v.5 (Andersen, et al. 2004) was used to evaluate the normalized count matrix to find the transcript with the highest stability value and lowest expression variation within and between the two conditions. The annotation and primer sequences of the genes transcripts used for qPCR are presented in supplementary table S8.

Total RNA for qPCR validation was extracted and purified as described above from larvae of both conditions. High-quality RNA (200 ng) was converted into cDNA using iScript cDNA synthesis kit (Bio-Rad). The specificity, dynamic range and PCR efficiency of each primer pair was determined by testing against a 6-step twofold dilution series of cDNA. All genes were amplified by triplicate (i.e. as technical replicates) in the two larvae from each condition (i.e. as biological replicates). All PCRs (20 ul final volume) were run on a CFX96 (Bio-Rad™) thermocycler using 300 nM of each primer, 10 ul iQ SYBR^®^ Green Supermix (Bio-Rad™) and 4ul of cDNA diluted 4-fold. PCR conditions used throughout were 95 °C for 3 min followed by 40 cycles of 95 °C for 10 s, 60 °C for 15 s and 72 °C for 10s. Melt curves were run after amplification to check for specificity from 55°C to 95°C with fluorescence readings taken in 0.5°C increments. Amplification efficiency of each gene was calculated by plotting the standard curve Cq values against the log of the dilution factor for each point on the curve. The relative change in gene expression between the treatments was examined using the delta Ct method (Schmittgen and Livak 2008).

### Unique Genes

I compared the transcriptomes from the two larval conditions in order to identify gene transcripts that are only expressed in one condition (i.e. unique gene transcripts). First, the Al_Trans and Col_Trans transcriptomes were aligned (BLASTn) against each other using one transcriptome at the time as the reference. Pairs of transcripts (i.e. from reference and query) longer than 1kb (± 10bp) that showed < 1% alignment score were considered as unique gene transcripts to either condition. The functional annotation of these transcripts was then obtained as above and the results graphically display in a tree map using R v3.3.3 (R Core Team 2017).

### Candidate Melanin Biosynthesis Genes

Here I took a candidate gene approach to evaluate expression patterns in genes from the melanin biosynthesis pathway in larvae from both conditions. The candidate genes investigated were tyrosine hydroxylase (*TH*), *yellow, laccase2*, DOPA decarboxylase (*Ddc*), arylalkylamine N-acetyltransferase (*aaNAT*), *tan*, and *ebony*. These genes or the enzymes they encode have been shown to impact insect black pigmentation and patterning (Fujii, et al. 2013; Liu, et al. 2016; van’t Hof and Saccheri 2010). In addition, I evaluate expression of tetrahydrobiopterin (BH4), a cofactor outside the melanin pathway that impacts the hydroxylation of tyrosine, the precursor of melanin synthesis (supplementary material S9). It was recently found that mutations in 6-pyruvoyl-tetrahydropterin synthase (*PTS*), the gene that encodes for BH4, can promote albinism in the silkmoth (Fujii, et al. 2013). Differences in expression (i.e. over-expression or down-regulation) in these candidate genes could be informative in understanding the albino condition. Lepidoptera mRNA sequences from each candidate gene were first obtained from NCBI. The sequences were then searched for orthology against the transcriptomes of both conditions through their protein translation to all six possible frames using tBLASTx, and blasted (BLASTx) back to NCBI to confirm orthology. Finally, the expression of the ortholog transcripts in each larval condition was evaluated using the TPM method described above and compared through a non-parametric t-test in R v3.3.3 (R Core Team 2017). The candidate gene transcript sequences and the orthologs accession numbers are given in the supplementary table S1.

## Results

### Transcriptome Assembly Validation and Annotation

A total of 32.5 million pair-end clean reads were obtained after trimming and filtering with an average of 15.4 (41% GC) and 12.2 (44% GC) million reads from the albino and coloured samples respectively. Overall, the two condition-specific transcriptomes were of comparable quality and completeness. The basic descriptive statistic of the transcriptomes is presented in table 1. Up to 56% and 62% of transcripts (i.e. unigenes, hereafter genes) from the Al_Trans and Col_Trans transcriptomes respectively, had functional annotation in the databases. Transcripts involved in nucleic acid metabolic process, ATP binding, and catalytic activity were more abundant in the Al_Trans, whereas genes in proteolysis, RNA-dependent DNA replication, translation, and endonuclease activity were more abundant in the Col_Trans (Figure 1). The full annotation results of the transcriptomes are presented in the supplementary tables S2 and S3.

**Table 1.** Descriptive statistics of the albino (Al_Trans) and coloured (Col_Trans) transcriptome assemblies of the wood tiger moth (*Arctia plantaginis*).

| Transcriptome | No.genes | N50 | Complete (%) | Single copy (%) | Fragmented (%) | Missing (%) |
| --- | --- | --- | --- | --- | --- | --- |
| Al_Trans | 28919 | 1105 | 67.0 | 58.6 | 15.6 | 4 |
| Col_Trans | 21787 | 1036 | 63.8 | 56.7 | 15.9 | 3 |

**Fig. 1.**
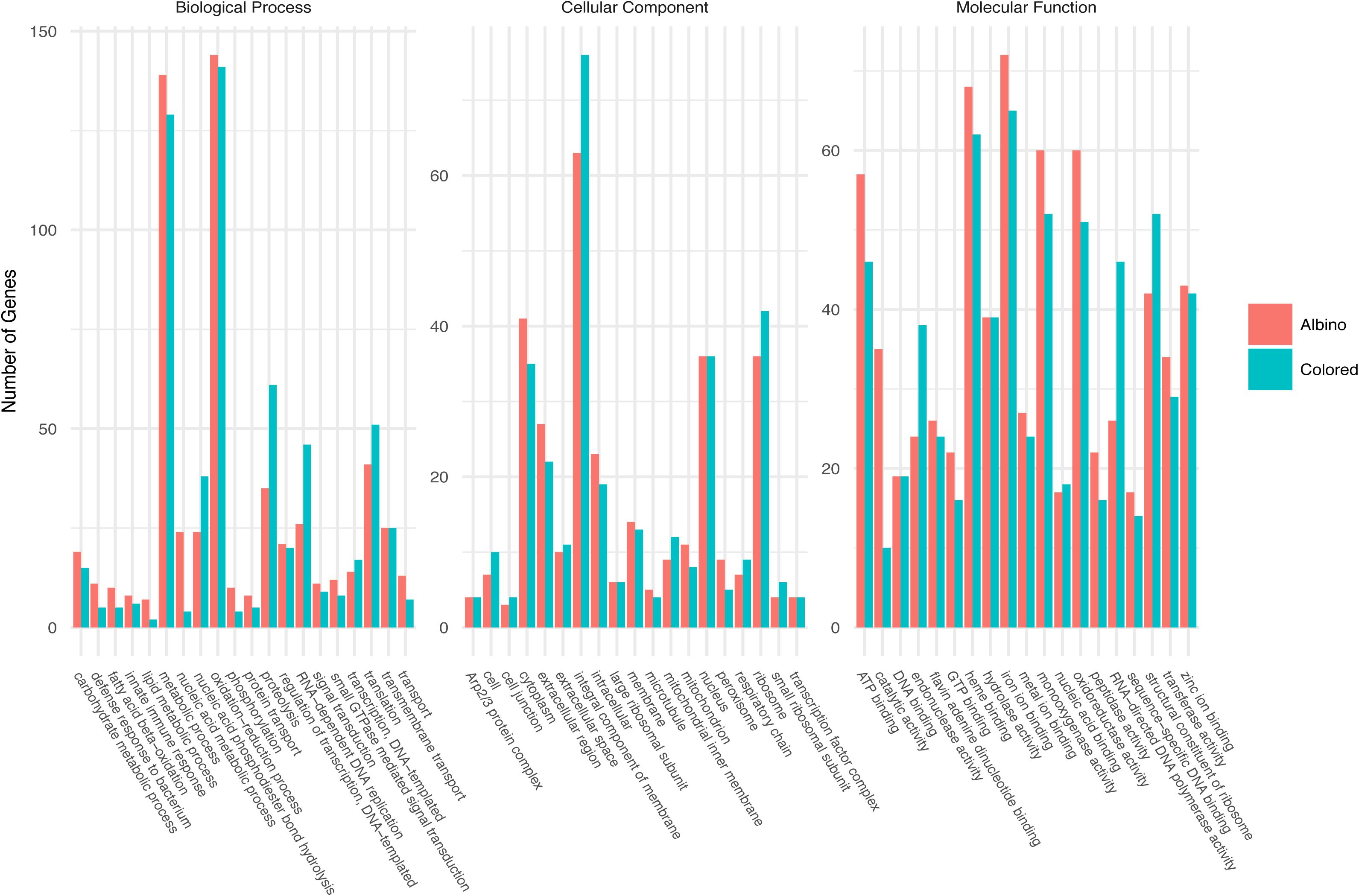
Functional annotation of expressed genes in albino and warningly coloured larvae of the wood tiger moth (*Arctia plantaginis*).

### Differential Gene Expression

A total of 302 genes were differentially expressed between the two larval conditions. Of these, 103 were up-regulated in the albino condition and 199 down-regulated in the coloured condition. The annotation analyses returned 57 up-regulated and 105 down-regulated genes with functional annotation (supplementary tables S4 and S5). Grouping of these genes by their functional annotation (GO terms) showed that immune system-defence response process, structural constituents of cuticle, and peptidase activity were the most down-regulated genes in the albino condition. In contrast, genes involved in metabolic processes and transmembrane transport were most abundant and up-regulated in the coloured condition (Figure 2). The qPCR validation of the candidate genes was in good agreement with the RNA-seq data. All genes examined showed the same pattern of up- or down-regulation in both data sets. Moreover, a significant correlation was observed (*r* = 0.94, *P* = 0.0046) between ΔCq form qPCR and the TPM values from RNA-seq (Figure 3).

**Fig. 2.**
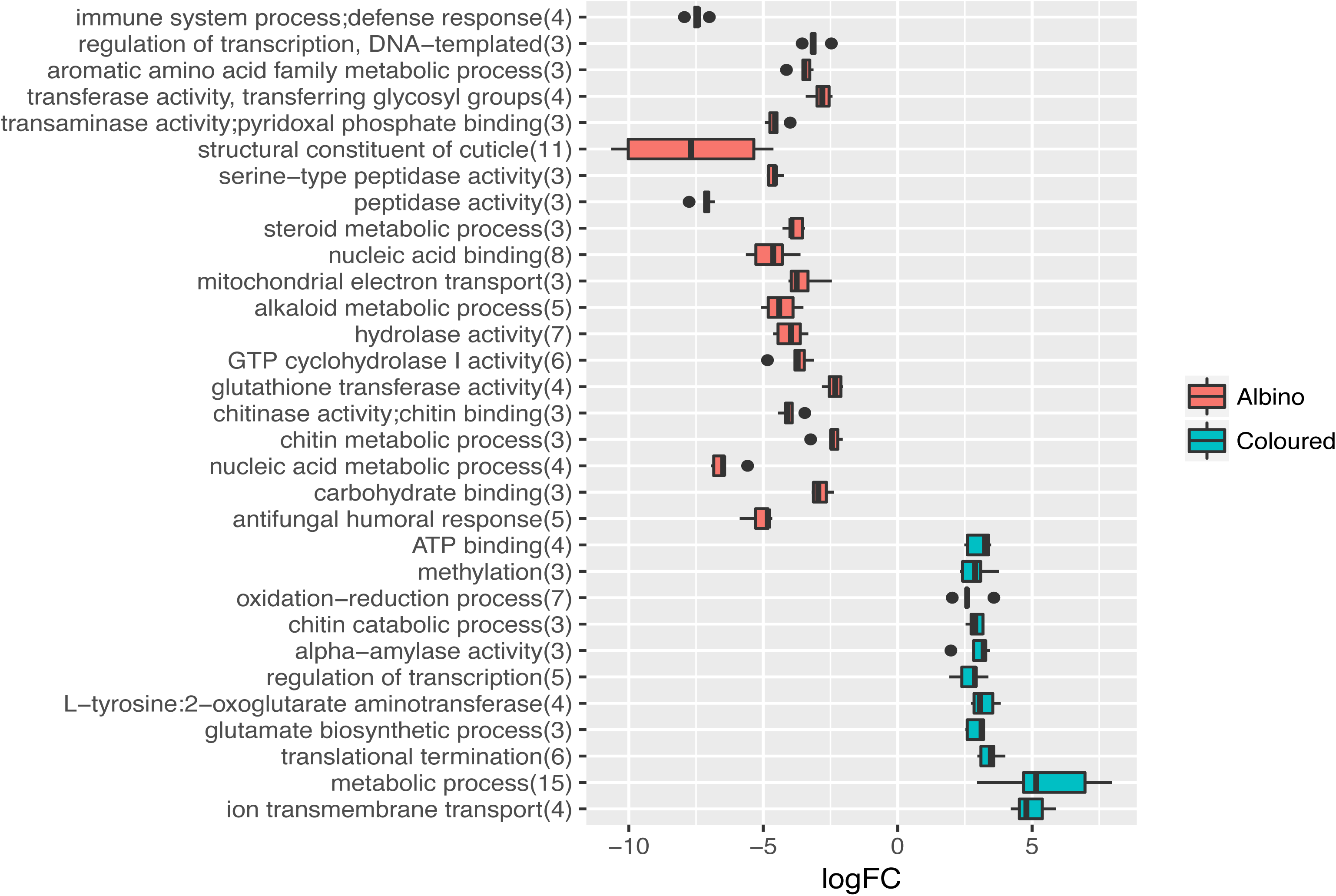
Differential gene expression between albino and coloured *Arctia plantaginis* larvae. y axis show biological, metabolic, and molecular functions. Numbers in parenthesis indicates the number of genes involved in those functions. x axis shows the fold chance in logarithmic scale.

**Fig. 3.**
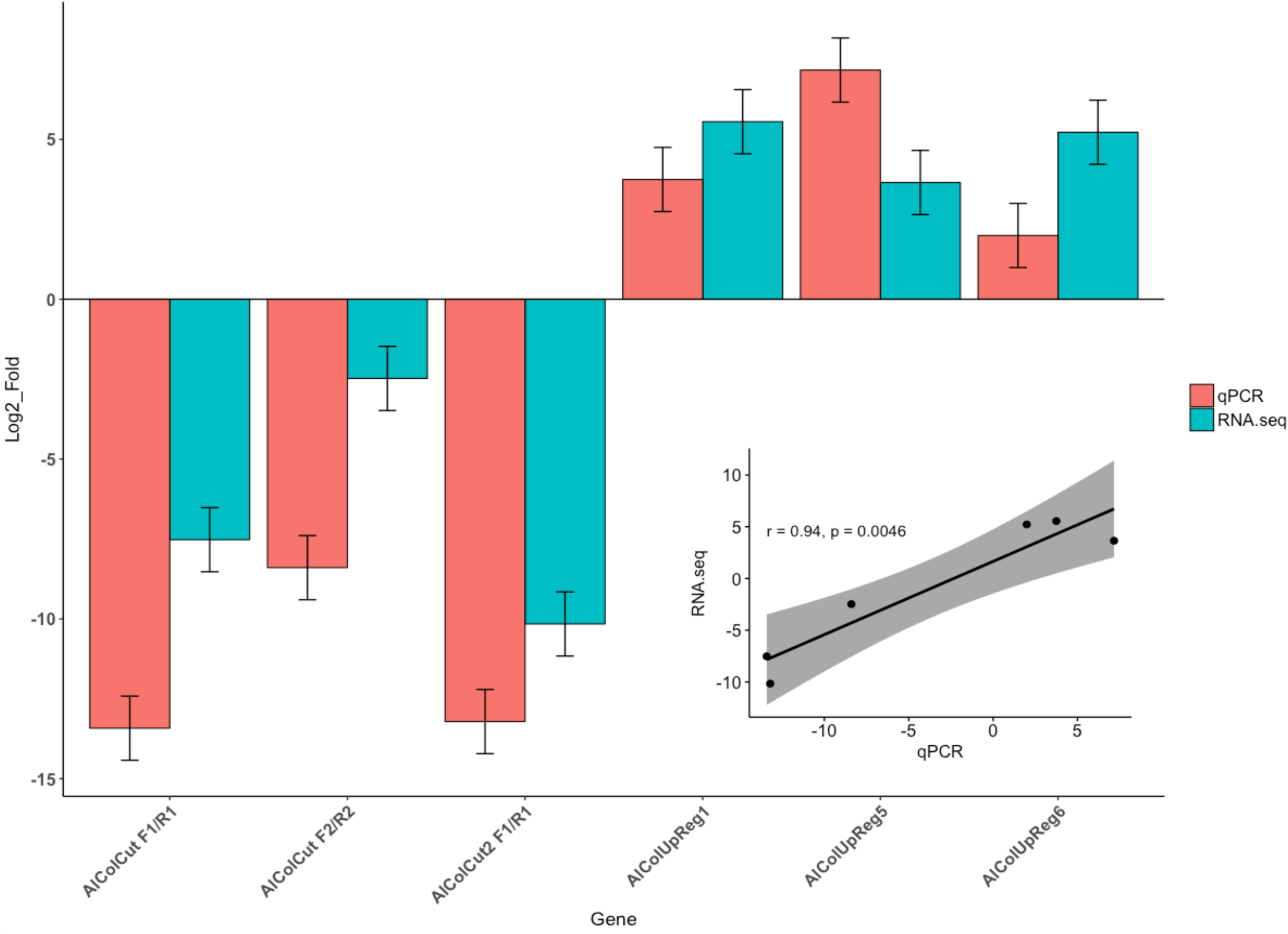
Log2 transformed Fold Change ΔCq values form qPCR and TPM from RNA-seq. Inset graph shows the correlation between datasets.

### Unique Genes

From the transcriptome-wide alignment comparison a total of 25 genes could be identified as unique to the Al_Trans, and 22 to the Col_Trans. Thirteen genes from the Al_Trans and 10 from the Col_Trans had functional annotation in public databases (supplementary tables S6 and S7). Gene transcripts involved in nucleotide biosynthesis, copper ion transmembrane transport, and nucleic acid binding were mostly represented in the Al_Trans (Figure 4), whereas intracellular protein transport, GTP binding, multicellular development and protein catabolism processes corresponded to the majority of unique genes in the Col_Trans (Figure 5). The full functional annotation is given in supplementary tables S6 and S7.

**Fig. 4.**
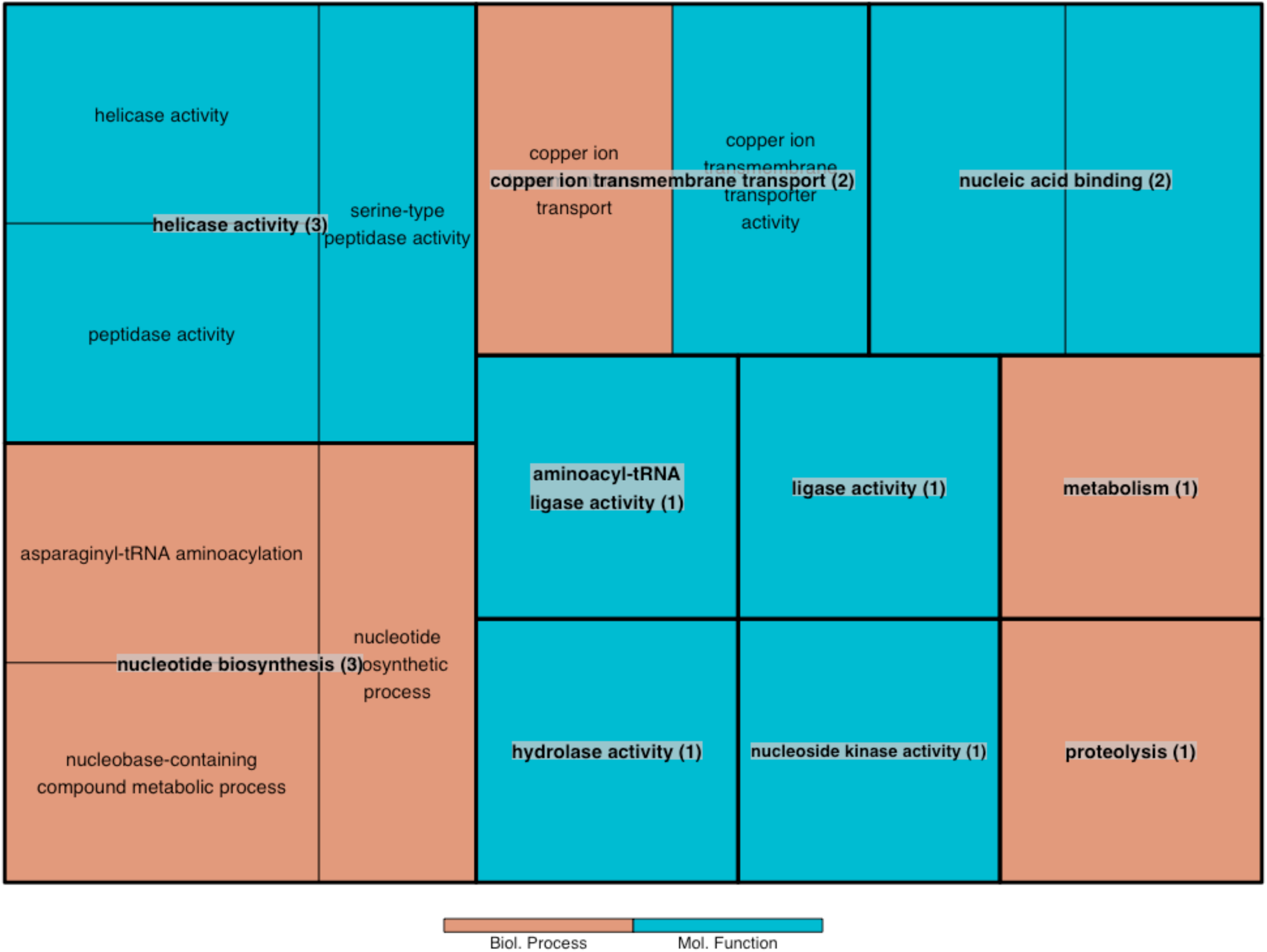
Biological processes and molecular functions of unique gene transcripts to the albino transcriptome (Al_Trans). Numbers in parenthesis indicate the number of genes observed in each process or function. The size of the squares is proportional to the number of genes involved in each category.

**Fig. 5.**
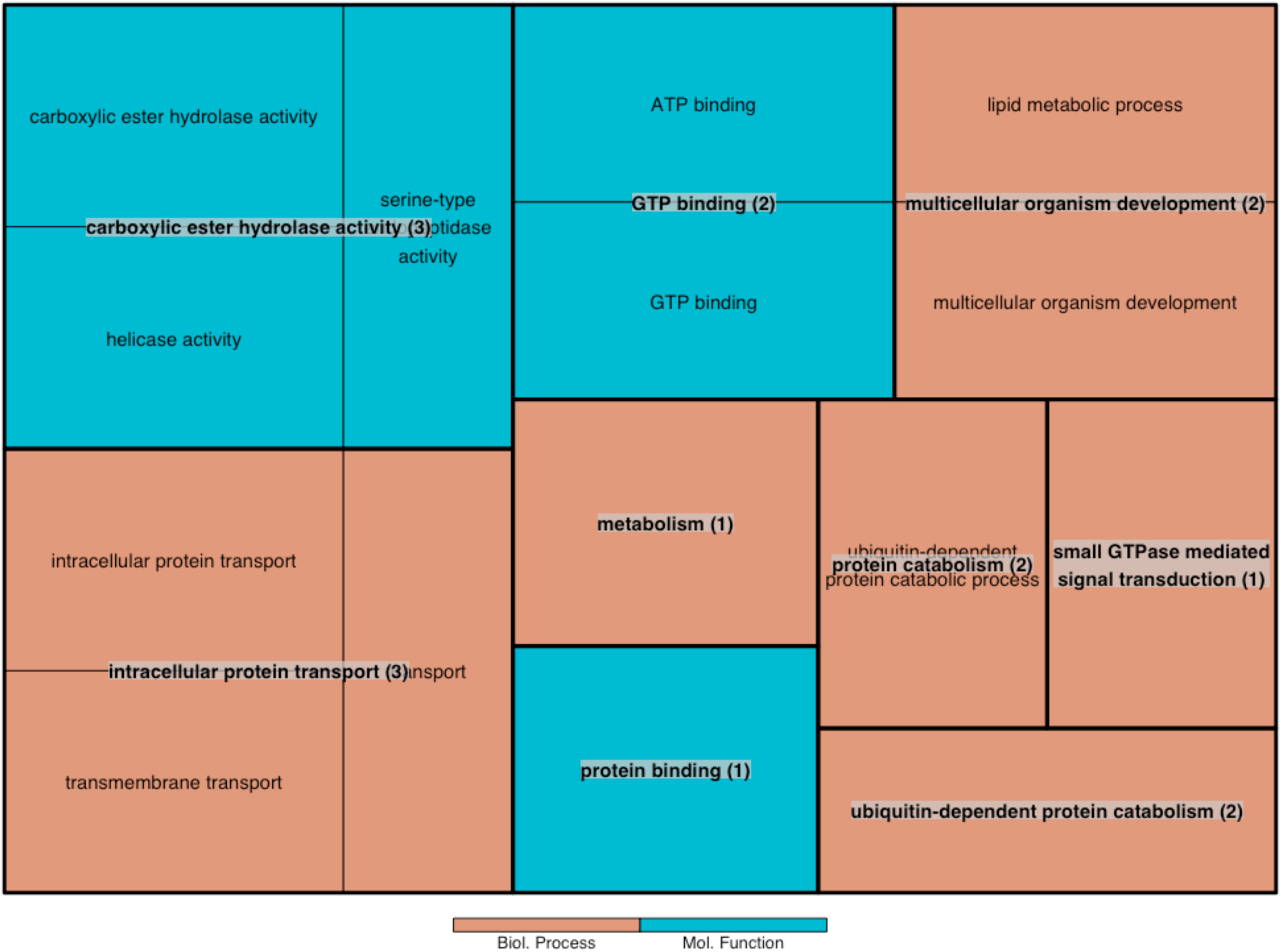
Biological processes and molecular functions of unique gene transcripts to the coloured transcriptome (Col_Trans). Numbers in parenthesis indicate the number of genes observed in each process or function. The size of the squares is proportional to the number of genes involved in each category.

### Candidate Melanin Biosynthesis Genes

Four candidate genes (*aaNAT, ebony, Ddc, TH*) showed significantly higher expression in the albino condition, whereas three others (*yellow, laccase2, tan*) were more expressed in the coloured condition. *PTS* on the other hand, showed non-significant differences (*P* > 0.05) in its expression levels between conditions (Table 2). The most differentially expressed genes were *TH* and *tan*, being highly expressed in the albino and coloured conditions respectively (Figure 6).

**Table 2.** Melanin biosynthesis genes t-test results between albino and coloured larvae of the wood tiger moth (*Arctia plantaginis*). The NCBI accession number of the candidate genes and the corresponding ortholog transcript ID from this study are shown. Cond shows if the gene was more expressed in the albino (Al) or coloured (Co) conditions.

| Gene | Accession No. | Transcript | Cond | P |
| --- | --- | --- | --- | --- |
| *aaNAT* | KP657626.1 | TRINITY_DN32379_c2_g2_i1 | Al | 0.023* |
| *tan* | KP657627.1 | TRINITY_DN36272_c0_g1_i2 | Co | 0.016* |
| *yellow* | AB525739.1 | TRINITY_DN24353_c1_g1_i1 | Co | 0.038* |
| *ebony* | AB439000.1 | TRINITY_DN37557_c4_g5_i1 | Al | 0.039* |
| *laccase2* | NM_001109925.1 | TRINITY_DN28256_c1_g1_i1 | Co | 0.032* |
| *Ddc* | GU953217.1 | TRINITY_DN22869_c0_g1_i1 | Al | 0.009* |
| *PTS* | ABD36088.1 | TRINITY_DN19160_c0_g1_i1 | Al | 0.097 |
| *TH* | GQ415559.1 | TRINITY_DN33765_c0_g1_i1 | Al | 0.005* |

**Fig. 6.**
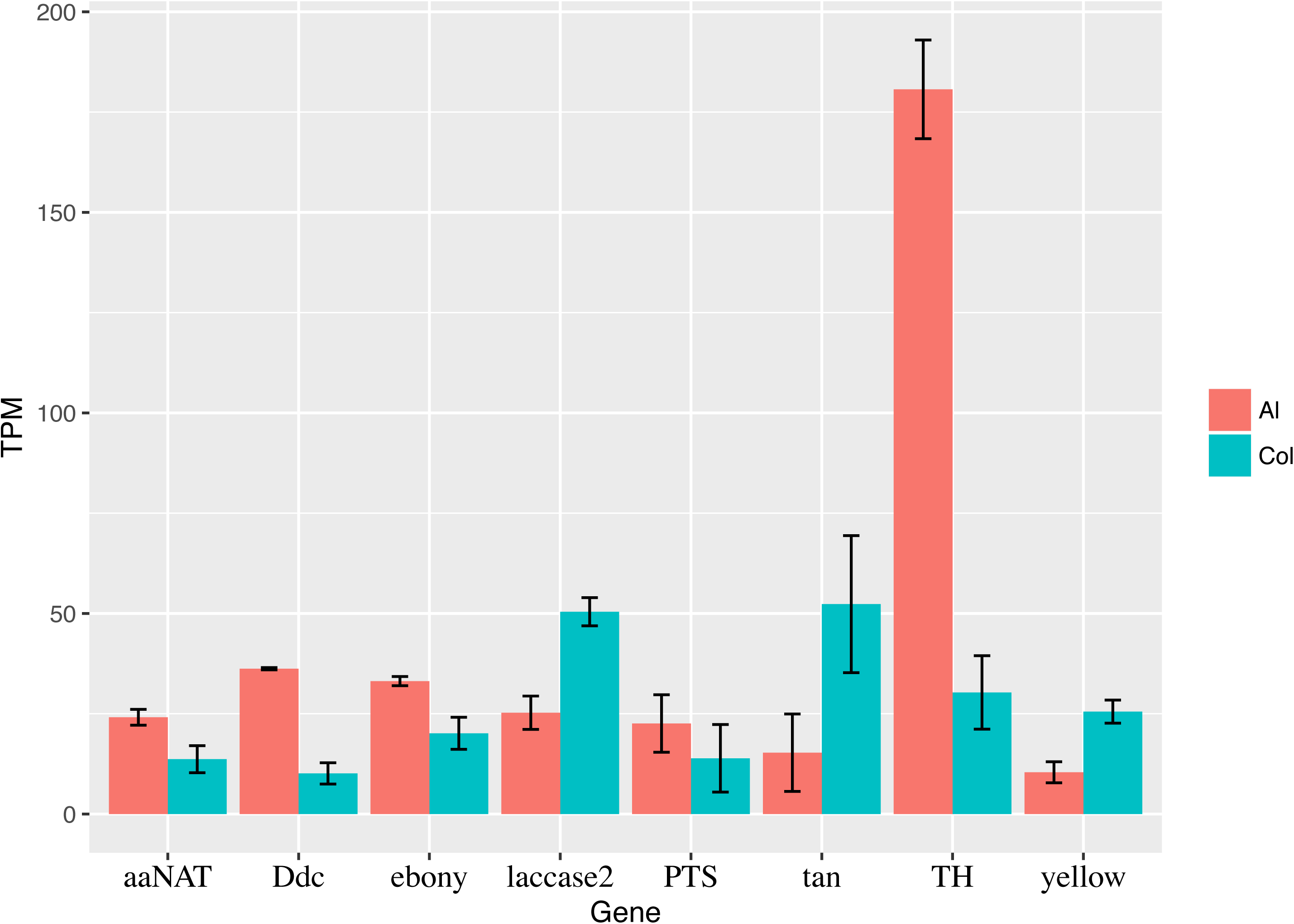
Candidate genes from the melanin biosynthesis pathway. Y-axis represents their expression level in transcripts observed per every million bases sequences (TPM). Error bars indicate the standard deviation between the expression of the two samples from each larval condition.

## Discussion

In this study, I report the transcriptome characterisation and comparison between albino wood tiger moth caterpillars and their warningly coloured siblings. The results showed substantial differences transcriptome-wide. Processes such as immune response and cuticle formation were found significantly reduced in albino caterpillars. Albinism also appears to be associated with depressed immunity and with a high activity in processes essential for genome replication and subsequent RNA transcription. Moreover, core-melanin genes (*TH, Ddc*) showed a higher expression in albino larvae. However, a down-regulation of genes (*yellow, laccase2*) involved in final melanin conversion was observed. Taken together, these results suggest that caterpillar albinism may not be due to a depletion of melanin precursor genes as it could be expected. In contrast, the albino condition may result from the combination of faulty melanin conversion late in its synthesis and structural deficiencies in the cuticle preventing its deposition.

### Albinism Frequency

Albinism in humans has been extensively studied and its frequency is well documented for distinct populations among different geographic regions (Montoliu, et al. 2014). For wild species however, little information is available about its prevalence. Albinism seems to be a common phenomenon in species living in lightless environments such as caves or subterranean habitats (Bilandžija, et al. 2012; Bilandžija, et al. 2017; Oliveira and Aguiar 2008; Protas, et al. 2006). However, it can be observed also in aboveground species. For instance, albino chorus frogs (*Pseudacris triseriata*) have been reported in frequencies of 7% and 12% during two consecutive years from natural ponds (Corn 1986). Likewise, scatter reports of albino Viperinae (*Vipera ammodytes, V. aspis, V. seoanei*, and *V. berus*) in Europe have been collected ranging from 1 to 16 observations depending on the species with an increase in frequency towards Nordic populations. (Krecsák 2008). In Lepidoptera, cases of partial albinism have been observed in alpine butterflies, *Erebia cpiphron silesiana* and *E.sudeiica sudetica* with a frequencies ranging from 0.03-1.4% in *E.cpiphron* to 0.7-3.9% in *E. sudeiica* (Kuras, et al. 2001). In the case of the wood tiger moth, no albinos have been observed in the wild. However, albino larvae from the rearing program are F1 form wild-caught parents, from which, one in every 5000 larvae displays the albino condition. This may serve as a lower-end proxy of albino frequencies in the wild, but it needs to be better established by field studies.

### Albinism and Predation

It can be envisaged that albinism would have mostly adverse effects on anti-predator strategies such as aposematism. However, this has not been examined in depth. In insects, there are only few predation experiments reported. Using four different wild-caught bird species (*Parus major, Parus caeruleus, Erithacus rubecula* and, *Sylvia atricapilla*) as predators, Exnerová et al (2006) found that white-black mutants of the aposematic firebug (*Pyrrhocoris apterus*) were more attacked that the red-black wild-type, and equally attacked as non-aposematic grey-black controls. On the other hand, in a later experiment, naïve, hand-reared *P.major* did not show any avoidance and attacked firebugs equally irrespective of colour (Svadová, et al. 2009). Data from a field study in two alpine butterflies (*E.cpiphron, E. sudeiica*) showed that predation marks and malformations were positively associate with albinism in both species (Kuras, et al. 2001). It is worth noting that these studies examined partial albinos having some degree of dark pigmentation. The effect of predation on wholly albinos like those reported here remains to be examined. As suggested by the previous studies (Exnerová, et al. 2006; Svadová, et al. 2009), an effective aposematic strategy relies in great part in predator learning and generalisation. It is unclear if albino larvae would benefit from the effect of novelty or if they are beyond the limits of predators’ scope of generalization. It is also unknown the effect of albinism regarding non-visual predators like ants and other invertebrate. More studies are needed to investigate this. However, this is a difficult task due to the rarity of albino individuals, particularly in wild populations. Nonetheless, given the very low frequencies of albinism it can be inferred that the albino condition has severe adverse effects in the survival/fitness of aposematic species.

#### The Importance of Dark Pigmentation

Dark pigmentation plays a fundamental role in aposematic insects as both dark and bright colours in a contrasting pattern are necessary to displaying an effective warning signal. In wood tiger moth larvae, the orange patch contrasts with the dark body conforming the warning signal, which is more effective against visual predators when the orange patch is large (Lindstedt, et al. 2009). The patch is made of clusters of chitin hairs pigmented with eumelanin and diet-derived flavonoids that give it its orange colouration (Lindstedt, et al. 2010). The role of the orange pigmentation in physiology is unknown, but it is likely negligible. The black hairs on the other hand, contain only eumelanin (Lindstedt, et al. 2016), a type of melanin that produces the black pigmentation. Melanin is central for insect immunity (Tsakas and Marmaras 2010). Experiments with high- and low-melanin wood tiger moth larvae have shown a better resistance to oral bacterial infections in high-melanin larvae (Zhang, et al. 2012). Likewise, more melanised larvae showed a faster encapsulation response to artificial implants than less melanised ones (Nokelainen, et al. 2013). By analogy, the complete absence of melanisation in albino larvae hints to a suppressed immunity. This notion is supported by the gene expression results showing a down-regulation of genes involved in immune system, defence response, and antifungal responses in the albino larvae (Figure 2). However, it has recently been shown that wound-healing melanisation can still occur in most albino cave-adapted adapted species, including insects (Bilandžija, et al. 2017). Whether this is the case in albino wood tiger moth is unknown. Future studies should investigate possible melanization responses to wounding or macroparasites in albinos from above ground species.

Melanic dark pigmentation not only correlates with insects’ immunity, but also protects from deleterious solar radiation, and at the same time helps in thermoregulation (Ellers and Boggs 2004; Stoehr and Goux 2008). It has been shown that darker wood tiger moth larvae are more efficient in thermoregulation than larvae with larger orange patches (Lindstedt, et al. 2009). More recently, it was found that darker larvae absorb more heat, keep higher body temperature, and actively avoid overheating by seeking shade sooner than less melanised larvae (Nielsen et al. in prep.). Such behavioural variation relative to the amount of dark pigmentation could be expected to be exaggerated in albino larvae. Accordingly, albinos may need partial sheltered thermoregulation due to the lack of protective dark pigmentation. This could result in longer heat-up periods, which in turn may reduce foraging time and also increase their exposure to non-visual predators. If true, albinos may face different predation pressures than their warningly coloured counterparts, for which, an aposematic strategy may be irrelevant.

Sclerotization is an independent but parallel process to melanisation used by insects for cuticle hardening and tanning. Freshly made cuticle is pale and soft, turning harder and sometimes darker within a few hours after moulting. Melanin is deposited into the cuticle via epidermal cells. The cuticle is composed of cross-linked chitin fibers and matrix protein forming an effective barrier against harmful influences from the environment such as desiccation, microorganisms, and predators (Andersen 2010). The cuticles of the albino wood tiger moths appeared softer and thinner than those of warningly coloured ones, suggesting deficient sclerotization, which may prevent melanin deposition. This is also indicated by the down-regulation of genes involved in chitin metabolic processes, structural constituents of cuticle, and chitinase activity and binding (Figure 2). Electron microscopy from albino larvae of the silkmoth (*Bombyx mori*) showed an abnormally loose cuticle structure compared to that of coloured ones, resulting in an inability to chew because of insufficient hardening of their mandibles (Tsujita and Sakurai 1971). Similar observations have been made in experimental albino mutants (via RNA interference) that showed a significant decrease in cuticle hardness (Gorman and Arakane 2010). Thus, structural deficiencies in albino cuticles may make them vulnerable to their surrounding environment, contributing to their low survival.

### The Melanin Biosynthesis Pathway

Previous research has indicated that insect pigmentation results from both regulatory and structural genes, with many of the structural genes encoding enzymes that are involved in the biochemical pathways that generate pigments. The genes and their biochemical transformations within the melanin synthesis pathway in Lepidoptera are highly conserved, and their spatio-temporal expression determines the location and abundance of pigmentation (Ferguson, et al. 2011a; Futahashi, et al. 2010; Yu, et al. 2011). Melanisation starts from the hydroxylation of tyrosine to 3,4-dihydroxyphenylalanine (DOPA) encoded by the gene *TH*. Phenol oxidase (PO) catalyzes the conversion of DOPA to Dopa-melanin encoded by the *yellow* together with *laccase2*, producing black melanin. In a parallel branch, DOPA can be transformed to dopamine through decarboxylation encoded by *Ddc* to produce dopamine-melanin, a brown melanin (supplementary material S9). It can be expected that albino larvae suffer from a suppression or depletion of these genes given their lack of melanic pigmentation. However, *TH* and *Ddc* showed higher expression in the albino larvae (Figure 6). Moreover, outside the melanin pathway, *PTS,* the cofactor impacting the melanin precursor *tyrosine* did not showed differences between the two larval conditions (Table 2, Figure 6). This indicates a non-depletion of master genes (*TH, Ddc, PTS*) in albino larvae, suggesting that melanin synthesis may be disrupted during the conversion of DOPA and Dopamine to melanin by *yellow* and *laccase2* genes. The observed low expression of these genes in albino larvae further supports this notion (Figure 6). Functional studies are needed to investigate biochemichal reactions and their impact in the melanin-promoting genes *lacasse2* and *yellow* family genes.

Downstream the melanin pathway, the production of dopamine-melanin can be suppressed by converting dopamine to *N*-b-alanyldopamine (NBAD). In this branch, β-alanine binds to dopamine by the activity of *ebony* forming NBAD, the precursor of yellow sclerotin. This can be reversed by the activity of *tan*, by which dopamine is converted back to dopamine melanin, thus promoting dark pigmentation. Finally, dopamine also can be converted to NADA sclerotin, which is colorless and encoded by *aaNAT* (supplementary material S9). Previous studies in Lepidoptera and other insects have found an up-regulation of *ebony* and *aaNAT* in non-black body regions, while *tan* is generally expressed in melanic regions (Ferguson, et al. 2011b; Futahashi, et al. 2008; Liu, et al. 2016; Osanai-Futahashi, et al. 2012; Zhan, et al. 2010). This is congruent with the results obtained here where the melanin-promoting genes *tan* and *yellow* showed a higher expression in the coloured larvae, whereas the melanin-suppressing genes *ebony* and *aaNAT* were more expressed in the albino larvae (Table2, Figure 6). Thus, in addition to a defective conversion of melanin from DOPA and Dopamine upstream the pathway, melanin may be further suppressed downstream the pathway in albino caterpillars.

## Conclusion

It is well recognized that neuro-sensorial, biochemical, metabolic and physiological anomalies are associated with the albinism condition in mammals (Creel 1980; Oetting and King 1999). Albino insects seem to suffer from similar pathological conditions. The albino caterpillars studied here showed a substantial differentiation in biological processes and metabolic functions compared to their warningly coloured full-sibs. Some symptoms can have severe deleterious effects in their physiology (i.e. supressed immunity) and in their structural properties (i.e. defective cuticle sclerotization). The occurrence of albinism in wild wood tiger moth populations and its effect on the aposematic strategy of the species are unknown. However, its frequency can be expected to be very low, and presumably, albinos experience different selection pressures due to behavioural differences induced by their peculiar physiology. More field surveys and experimental studies are needed to explore these notions.

## Acknowledgements

This study was funded by the Academy of Finland via the Centre of Excellence in Biological Interactions. I would like to thank Kaisa Suisto and greenhouse interns for spotting albino larvae. Atsushi Honma for providing photographs of albino larvae. Sari Viinikainen and Kishor Dhaygude for laboratory and analyses assistance. Carita Lindstedt for helpful discussions in improving this manuscript.

